# Decreased green autofluorescence intensity of lung parenchyma is a potential non-invasive diagnostic biomarker for lung cancer

**DOI:** 10.1101/343533

**Authors:** Mingchao Zhang, Yujia Li, Jie Zhang, Haohua Teng, Qing Zhang, Zhenzhen Xiang, Qing Chang, Yue Tao, Tianqing Chu, Weihai Ying

**Author notes:** Corresponding authors: Weihai Ying, Ph.D., Professor, School of Biomedical Engineering and Med-X Research Institute, Shanghai Jiao Tong University, Shanghai, 200030, P.R. China, Tianqing Chu, M.D., Ph.D., Associate Chief Physician, Department of Pulmonary Medicine, Shanghai Chest Hospital, Shanghai Jiao Tong University, Shanghai, 200030, P.R. China.

## Abstract

Early and non-invasive diagnosis is critical for enhancing the survival rates of lung cancer. In current study we determined the green autoflorescence (AF) of the pulmonary parenchyma of lung cancer patients, using 488 nm and 500 - 550 nm as excitation wavelength and emission wavelength, respectively. Our study has suggested that decreased green AF intensity of pulmonary parenchyma may be a potential diagnostic biomarker for lung cancer: First, the green AF intensity of the cancerous tissues is less than 40% of those of distant non-neoplastic tissues and peri-neoplastic tissues; second, the green AF intensity of both the distant non-neoplastic tissues and cancerous tissues of squamous carcinoma is significantly lower than that of adenocarcinoma; third, the AF intensity of the peri-neoplastic tissues of the lung cancer patients is negatively correlated with the stages of lung cancer; and fourth, our study on the AF spectrum of pulmonary parenchyma has suggested that the AF may result from the AF of the keratins or FAD of the pulmonary parenchyma. Collectively, our study has suggested that decreased green AF intensity of pulmonary parenchyma may become a novel biomarker for non-invasive diagnosis of lung cancer. The green AF intensity may also be used to non-invasively differentiate squamous carcinoma and adenocarcinoma, as well as the stages of lung cancer. Moreover, our diagnostic approach for lung cancer may be used for image-guided surgery of lung cancer for lesions that have not been identified by CT.

## Introduction

Lung cancer is the leading cause of cancer deaths globally (11). Approximately 15% of lung cancer is small cell lung cancer (SCLC), while approximately 85% of lung cancer is non-small cell lung cancer (NSCLC) (3). Adenocarcinoma and squamous cell carcinoma are two major histological subtypes of NSCLC (3). Patients diagnosed with distant metastatic disease (stage IV) had a 1-year survival rate of 15-19%, while surgical resection of NSCLC offers a favorable prognosis, with 5-year survival rates of 70 – 90% for small, localized tumors (stage I) (4,9). However, at the time of diagnosis, approximately 75% patients around the world have advanced disease (stage III/IV) (12).

The major problems in current diagnostic approaches for lung cancer include: 1) CT-based diagnosis of lung cancer is of low precision, while the precision rate of PET/CT in diagnosis of the pulmonary modules smaller than 1 cm^2^ is only approximately 30%; 2) electronic bronchoscope can reach only subsegmental bronchus, which can obtain samples only from the lesions along bronchial pathway for pathological analyses; 3) Fluorescence Bronchoscopy can be used only for the cancer diagnosis of Trachea mucous membrane above segment, while it can not provide cancer diagnosis on subsegmental bronchial mucosa; and 4) another widely used invasive diagnostic approach for lung cancer - Aspiration biopsy of pulmonary nodules – has several limitations, including its low capacity in localizing pulmonary modules smaller than 1 cm^2^ and its low capacity to obtain sufficient number of cells from the lesions for diagnosis. Collectively, due to the limitations of the current diagnostic approaches for lung cancer, numerous patients without lung cancer have to undergo unnecessary surgery. The low capacity in providing precise diagnosis before surgery also significantly increases the challenges for the surgeons to make correct judgment during surgery. Moreover, so far there has been no approach to discover the lung cancer lesions that have not been identified by CT during surgery. Therefore, novel, non-invasive and efficient diagnostic approaches for lung cancer are critically needed.

Skin autofluorescence (AF) has shown promise for non-invasive diagnosis of such diseases as diabetes (6,7). Our recent study has found that UV-induced epidermal green AF, which is originated from UV-induced keratin 1 proteolysis, can be used as a new biomarker for predicting UV-induced skin damage (5). Since it is non-invasive and simple to detect AF, it is of significance to search for the potential of AF as new biomarkers for diseases.

In current study we determined the green autoflorescence (AF) of the distant non-neoplastic tissues, peri-neoplastic tissues and cancerous tissues of the pulmonary parenchyma of lung cancer patients, using 488 nm and 500 – 550 nm as excitation wavelength and emission wavelength, respectively. Our study has suggested that decreased green AF intensity may become a novel potential diagnostic biomarker for lung cancer:

## Methods

### Human subjects

This study was conducted according to a protocol approved by the Ethics Committee of Shanghai Chest Hospital, Shanghai Jiao Tong University. The lung cancer patients included adenocarcinoma and squamous-cell carcinoma patients, who were hospitalized in the Department of Pulmonary Medicine, Shanghai Chest Hospital, Shanghai Jiao Tong University. The age of the patients is 63.15 ± 5.46 years of old.

### Autofluorescence determinations

The green AF of the tissues were photographed under a two-photon fluorescence microscope. The excitation wavelength is 485 nm, and the emission wavelength is 500 – 550 nm.

### Animal studies

Male C57BL/6Slac mice of SPF grade were purchased from SLRC Laboratory (Shanghai, China). All of the animal protocols were approved by the Animal Study Committee of the School of Biomedical Engineering, Shanghai Jiao Tong University.

### Determinations of the AF spectrum of pulmonary parenchyma of mice

C57BL/6Slac mice were anesthetized by intraperitoneal injection of 3% bromethol. The spectra of the AF of the pulmonary parenchyma of the mice was determined by using a two-photon fluorescence microscope (A1 plus, Nikon Instech Co., Ltd., Tokyo, Japan). The excitation wavelengths were 700 nm, 760 nm, 800 nm, and 850 nm. After the imaging, the images were analyzed automatically.

### Statistical analyses

All data are presented as mean ± SEM. Data were assessed by one-way ANOVA, followed by Student - Newman - Keuls *post hoc* test, except where noted. *P* values less than 0.05 were considered statistically significant.

## Results

### 1) The green AF of the cancerous tissues is less than 40% of those of distant non-neoplastic tissues and peri-neoplastic tissues

Using 488 nm and 500 – 550 nm as excitation wavelength and emission wavelength, respectively, we determined the AF of the cancerous tissues, peri-neoplastic tissues and distant non-neoplastic tissues of 38 lung cancer patients. Our study showed that the green AF of the cancerous tissues was obviously lower than those of distant non-neoplastic tissues and peri-neoplastic tissues (Fig. 1A). Quantifications of the AF images indicated that the green AF of the cancerous tissues is less than 40% of those of distant non-neoplastic tissues and peri-neoplastic tissues (Fig. 1B).

**Fig. 1.**
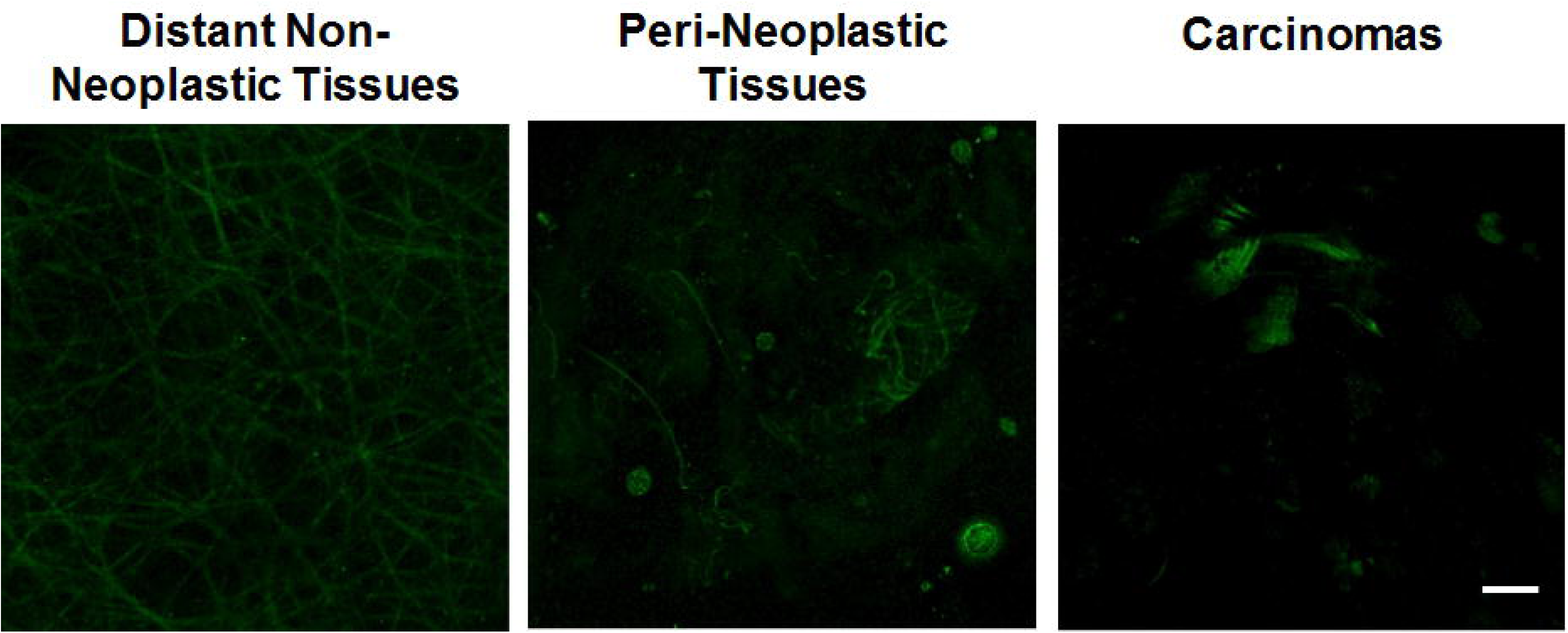
The green AF of the cancerous tissues is less than 40% of those of distant non-neoplastic tissues and peri-neoplastic tissues. (A) Using 488 nm and 500 – 550 nm as excitation wavelength and emission wavelength, respectively, we determined the AF of the cancerous tissues, peri-neoplastic tissues and distant non-neoplastic tissues of 38 lung cancer patients. Our study showed that the green AF of the cancerous tissues was obviously lower than those of distant non-neoplastic tissues and peri-neoplastic tissues. (B) Quantifications of the AF images have indicated that the green AF of the cancerous tissues is less than 40% of those of distant non-neoplastic tissues and peri-neoplastic tissues. N = 30. **, *p* < 0.01; ***, *p* < 0.001.

### 2) The green AF of both the distant non-neoplastic tissues and cancerous tissues of squamous carcinoma patients is significantly lower than that of adenocarcinoma patients

We also compared the green AF of the tissues of squamous carcinoma patients with that of adenocarcinoma patients. For both the squamous carcinoma patients and the adenocarcinoma patients, the green AF of the cancerous tissues was significantly lower than that of the distant non-neoplastic tissues and peri-neoplastic tissues (Fig. 2). However, the AF of both the distant non-neoplastic tissues and the cancerous tissues of the squamous carcinoma patients was significantly lower than that of the adenocarcinoma patients (Fig. 2).

**Fig. 2.**
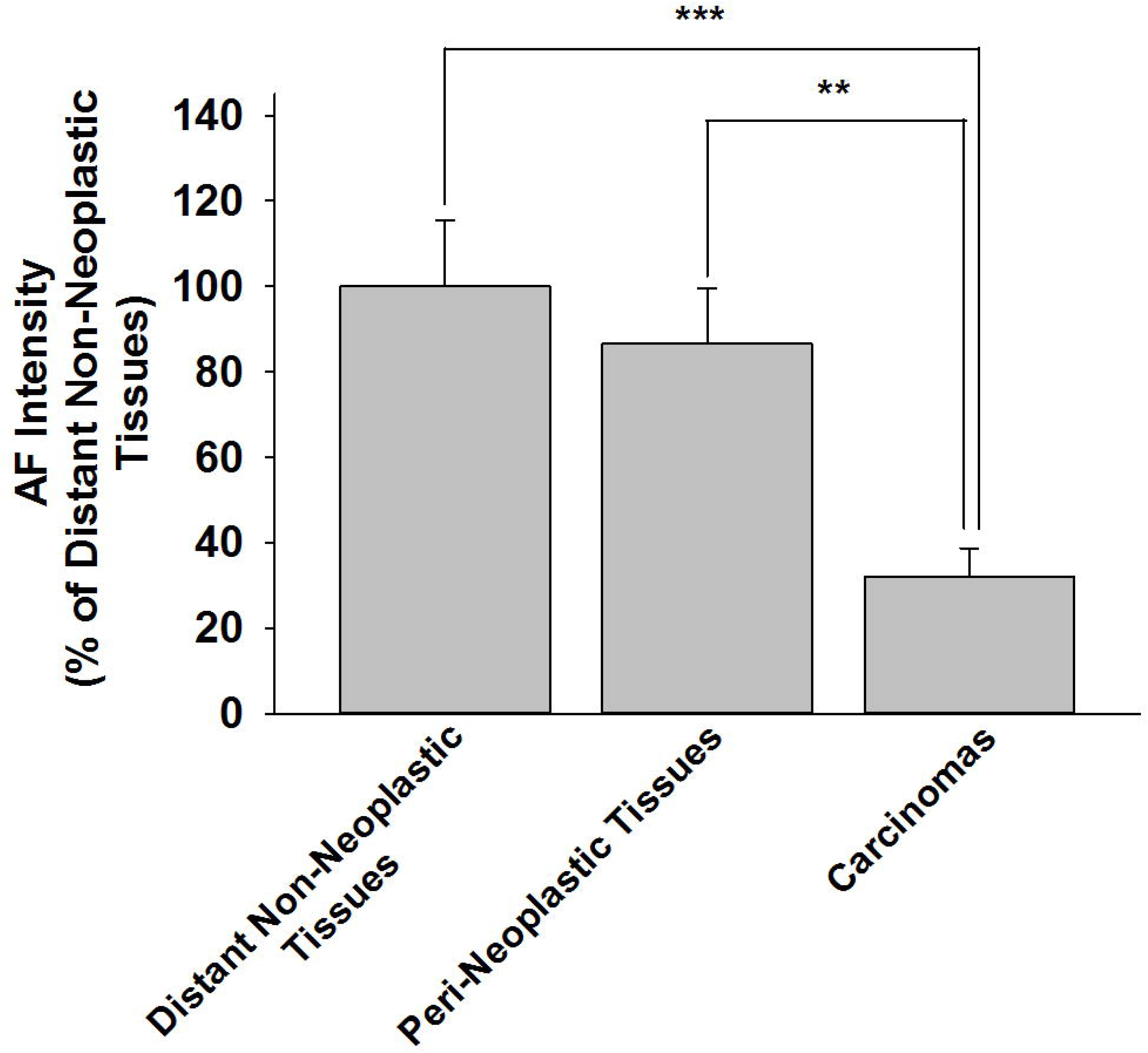
The green AF of both the distant non-neoplastic tissues and cancerous tissues of squamous carcinoma is significantly lower than that of adenocarcinoma patients. We compared the green AF of the tissues of squamous carcinoma patients and that of adenocarcinoma patients. For both the squamous carcinoma patients and the adenocarcinoma patients, the green AF of the cancerous tissues was significantly lower than that of the distant non-neoplastic tissues and peri-neoplastic tissues. However, the AF of both the distant non-neoplastic tissues and the cancerous tissues of the squamous carcinoma patients was significantly lower than that of the adenocarcinoma patients. N = 11 - 16. *, *p* < 0.05; **, *p* < 0.01; ***, *p* < 0.001; #, *p* < 0.05 (Student *t*-test).

### 3) The AF intensity of the peri-neoplastic tissues of the lung cancer patients is negatively correlated with the stages of lung cancer

We also determined the green AF of the lung cancer patients at various stages. There was a strong negative correlation between the AF intensity of the peri-neoplastic tissues of the lung cancer patients and the stages of lung cancer (R^2^ = 0.98) (Fig. 3). In contrast, in both cancerous tissues and distant non-neoplastic tissues, there was no significant correlation between the AF intensity of the lung cancer patients and the stages of lung cancer (Fig. 3).

**Fig. 3.**
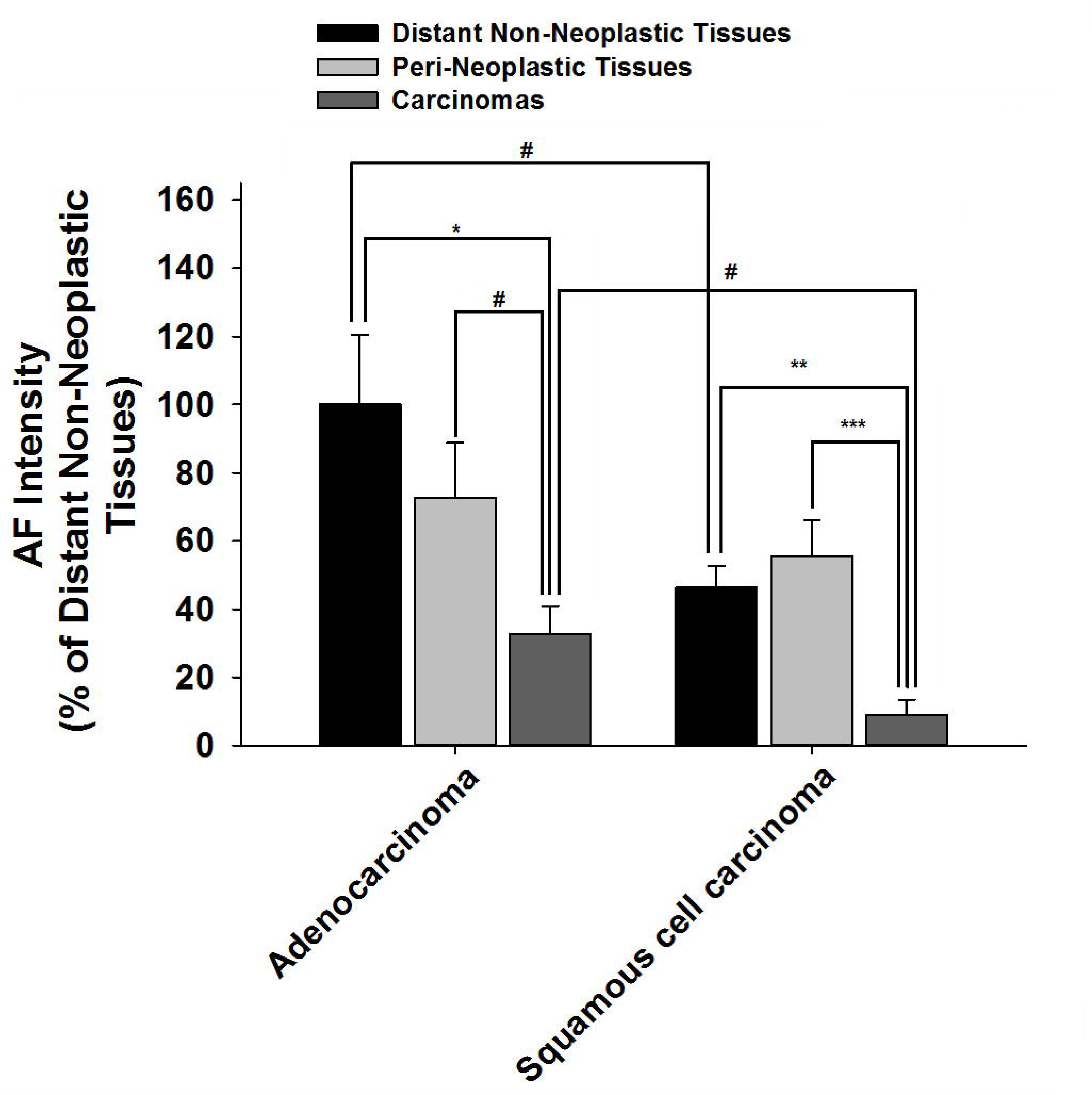
The AF intensity of the peri-neoplastic tissues of the lung cancer patients is negatively correlated with the stages of lung cancer. We also determined the green AF of the lung cancer patients at various stages. There was strong negative correlation between the AF intensity of the peri-neoplastic tissues of the lung cancer patients and the stages of lung cancer (R^2^ = 0.98). In contrast, in both cancerous tissues and distant non-neoplastic tissues, there was no significant correlation between the AF intensity of the lung cancer patients and the stages of lung cancer. The number of the patients in the stage of IA – IIA, IIB, IIIA, and IIIB is 10, 6, 7, and 3, respectively.

### 4) The study on the AF spectrum of the pulmonary parenchyma of mice has suggested that the AF may result from the keratins or FAD of the pulmonary parenchyma

We used two-photon fluorescence microscope to determine the spectra of the AF of the left upper lobe, left inferior lobe, right upper lobe, right middle lobe and right inferior lobe of the pulmonary parenchyma of mice. At the excitation wavelength of 760 nm, all of these spectra appeared to be highly similar to each other (Fig. 4A). The spectra match that of keratins and FAD, suggesting that the AF may result from the keratins or FAD of the pulmonary parenchyma. The AF intensity of left upper lobe was also determined when excitation wavelengths were 700, 760, 800 and 900 nm, respectively (Fig. 4B).

**Fig. 4.**
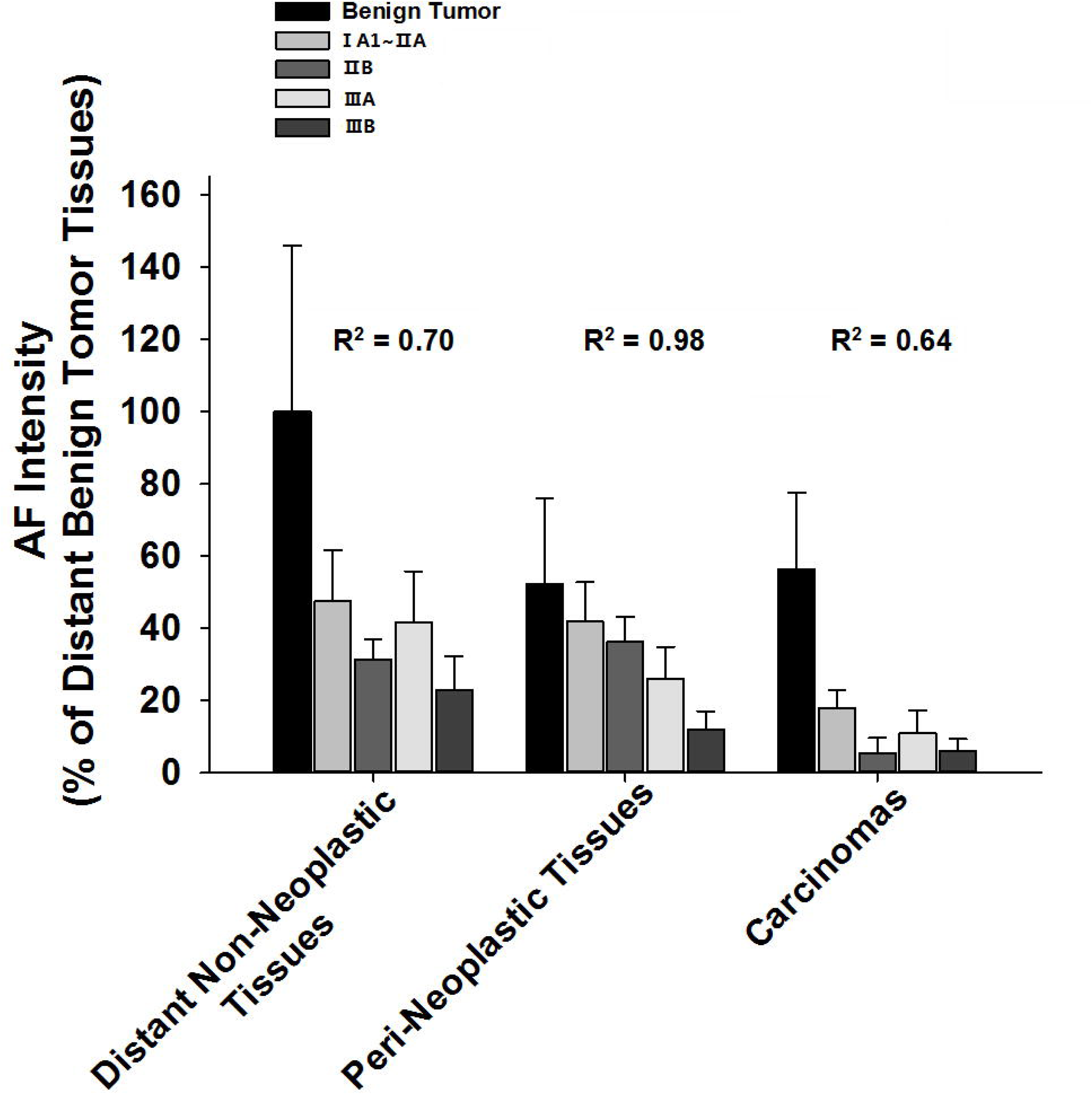
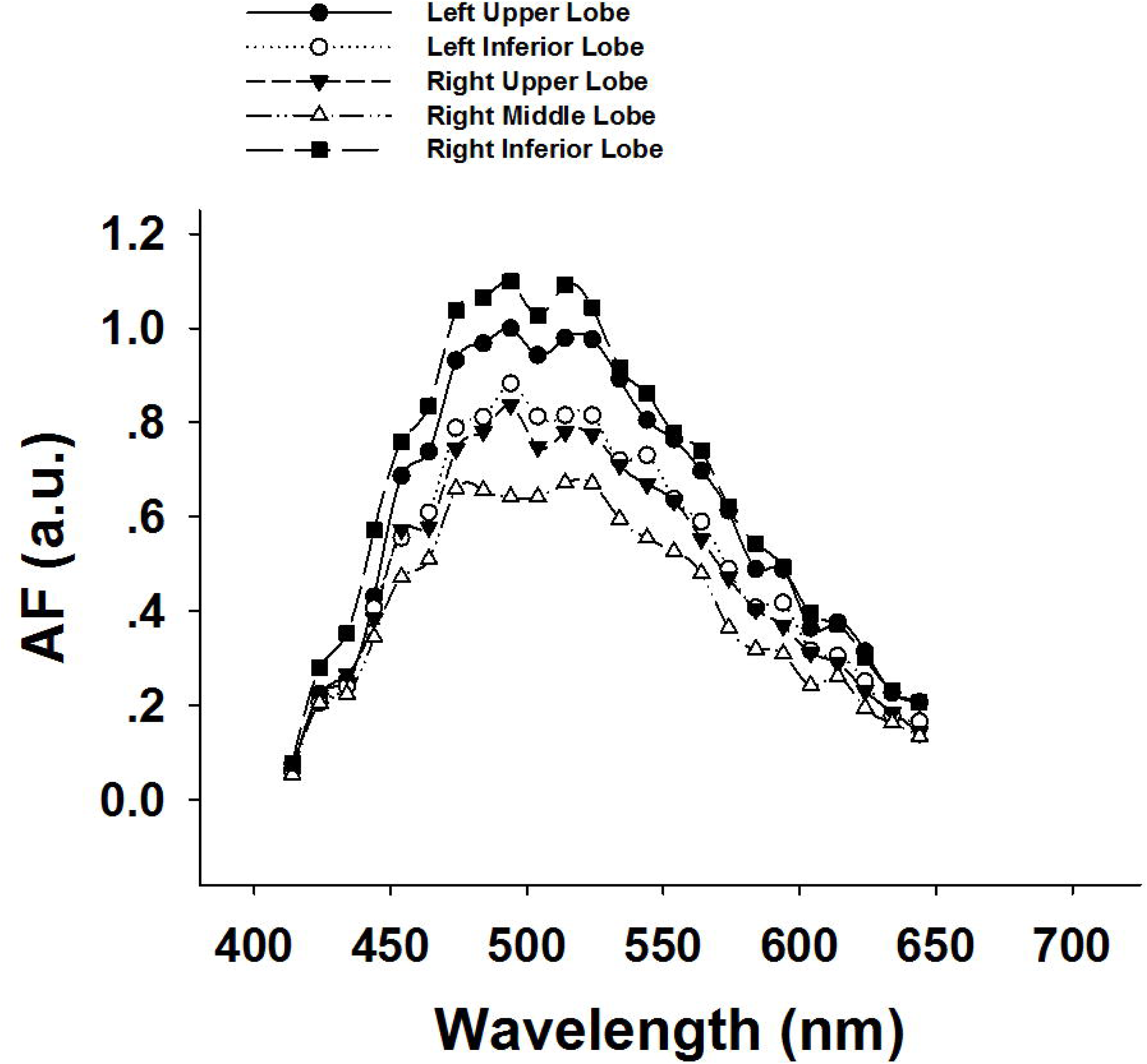
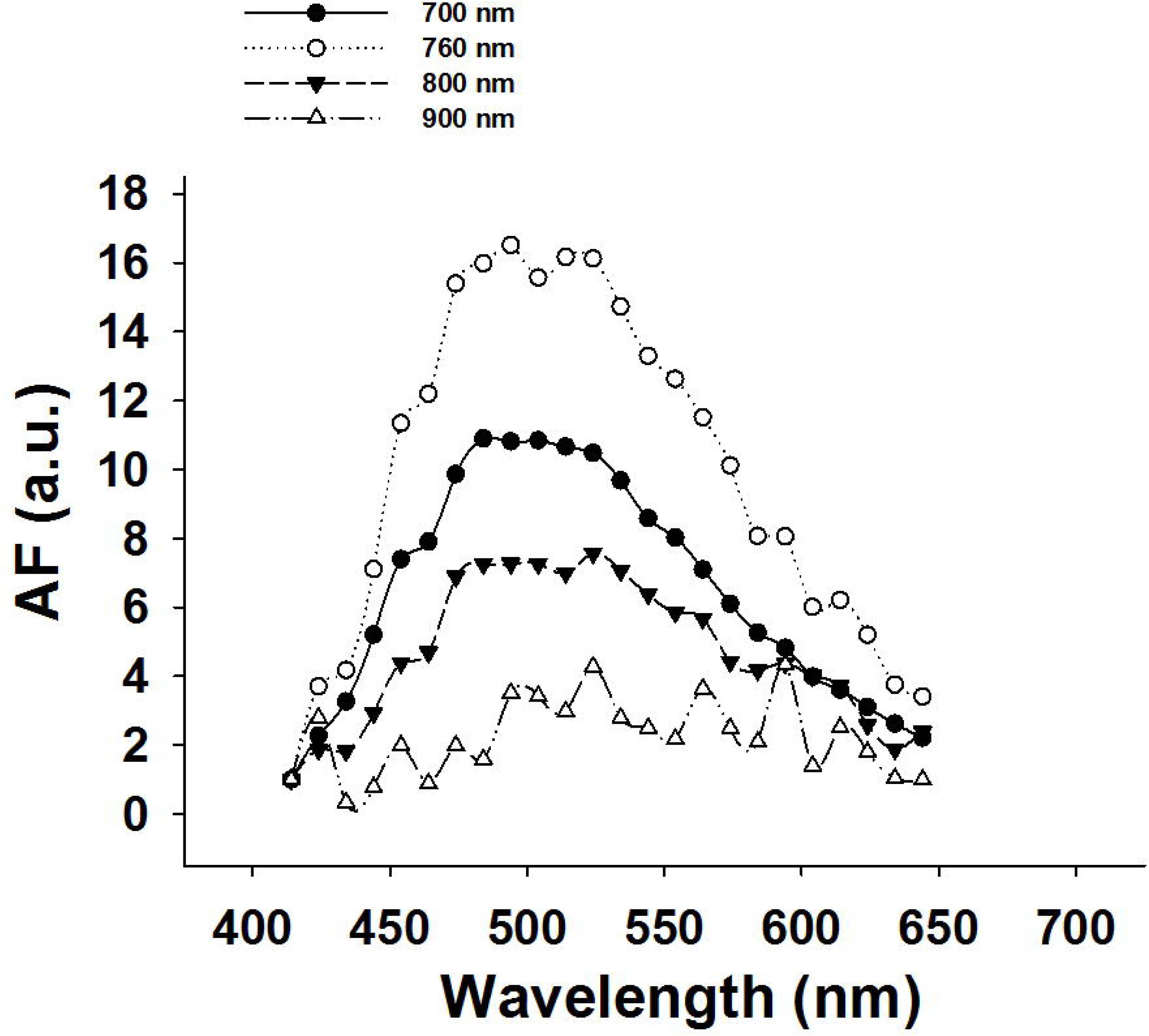
The study on the AF spectrum of the pulmonary parenchyma of mice has suggested that the AF may result from the keratins or FAD of the pulmonary parenchyma. (A) We used two-photon fluorescence microscope to determine the spectra of the AF of the left upper lobe, left inferior lobe, right upper lobe, right middle lobe and right inferior lobe of the pulmonary parenchyma of mice. At the excitation wavelength of 760 nm, all of these spectra appeared to be highly similar to each other. N = 3. (B) The AF intensity of left upper lobe was also determined when excitation wavelengths were 700, 760, 800 and 900 nm, respectively. N = 3.

## Discussion

The major findings of our study include: First, the green AF of the cancerous tissues is less than 40% of those of distant non-neoplastic tissues and peri-neoplastic tissues; second, the green AF of both the distant non-neoplastic tissues and cancerous tissues of squamous carcinoma is significantly lower than that of adenocarcinoma patients; third, the AF intensity of the peri-neoplastic tissues of the lung cancer patients is negatively correlated with the stages of lung cancer; and fourth, our study on the AF spectrum has suggested that the AF may result from the keratins or FAD of the pulmonary parenchyma.

Our study has suggested a novel approach for non-invasive diagnosis of lung cancer by determining the AF intensity with excitation wavelength and emission wavelength of 488 nm and 500 – 550 nm, respectively: The cancerous lesions have profoundly lower AF intensity compared with non-neoplastic tissues and peri-neoplastic tissues. This new non-invasive approach may replace the currently widely used invasive approach for lung cancer diagnosis - the aspiration biopsy of pulmonary nodules: The green AF intensity of the pulmonary nodules that have been identified by CT as well as the green AF intensity of multiple other regions in the pulmonary parenchyma should be determined by an AF imaging device, which can be designed as a new type of Fluorescence Bronchoscope or an micro-device of imaging linked with a micro-needle. The pulmonary nodules may be diagnosed as cancerous tissues if the modules have significantly lower green AF intensity compared with that of the other regions of the pulmonary parenchym.

It is important to elucidate the mechanisms underlying the decreases in the green AF intensity in lung cancer. There are multiple types of keratins in the lung tissues (2,13), while both keratins and FAD are major known fluorophores (1,8). Our study has shown that the AF spectrum of lung tissues matches that of keratins and FAD (10,14). We propose that the profoundly decreased green AF intensity of lung cancer tissues may result from significant loss of keratins and/or FAD of the lung cancer tissues. Future studies are warranted to further test the validity of our proposal.

Our study has also suggested that the **g**reen AF of both the distant non-neoplastic tissues and cancerous tissues of squamous carcinoma is significantly lower than that of adenocarcinoma patients. This observation has suggested that our AF-based approach may also be used to provide a preliminary judgment for differentiating squamous carcinoma and adenocarcinoma: If a pulmonary nodule has particularly low intensity of green AF, there is a high probability that the lung cancer is squamous carcinoma. In contrast, if a pulmonary nodule has mildly low intensity of green AF, there is a high probability that the lung cancer is adenocarcinoma. Future studies are needed to establish strong data base for enhance the precision to differentiate squamous carcinoma and adenocarcinoma.

It is expected that with future applications of artificial intelligence technology and big data technology, our green AF-based diagnostic approach for lung cancer may have significant increases in sensitivity and specificity. Based on our observations regarding the profoundly decreased green AF intensity, which probably results from decreased AF of keratins and/or FAD, we propose that decreased levels of FAD and/or certain isoforms of keratins may be key pathological changes of lung cancer.

Moreover, our diagnostic approach for lung cancer may also be used for image-guided surgery of lung cancer for the lesions that have not been identified by CT: The lung tissues that have exceedingly low green AF intensity may be the cancerous lesions, which could be removed by surgery. Future development of our diagnostic approach may profoundly enhance our capacity to provide precise guidance for lung cancer surgery to remove the cancerous lesions that have not been identified by CT.

## Acknowledgment

The authors would like to acknowledge the financial support by Major Research Grants from the Scientific Committee of Shanghai Municipality #16JC1400500 and #16JC1400502 (to W.Y.) and Chinese National Natural Science Foundation Grant #81271305 (to W. Y.).

